# Supervised Domain Adaptation Mitigates Cross-Ethnicity Prediction Error in Neuroimaging-Based Cognitive Prediction

**DOI:** 10.64898/2026.05.25.727742

**Authors:** Farzane Lal Khakpoor, William van der Vliet, Jeremiah Deng, Yue Wang, Narun Pat

**Affiliations:** Department of Psychology, University of Otago, Dunedin, New Zealand; School of Computing, University of Otago, Dunedin, New Zealand

**Author notes:** Corresponding Author: Narun Pat, PhD, Department of Psychology, University of Otago, William James Building, 275 Leith Walk, Dunedin 9016, New Zealand.

**Keywords:** Domain Adaptation, Neuroimaging, Machine Learning Fairness, Predictive Modelling, Sample Imbalance

## Abstract

Machine-learning models are increasingly used to predict cognitive and clinical outcomes from neuroimaging data, yet challenges in fairness and generalizability remain. Large-scale datasets are often racially and ethnically imbalanced, leading to systematic performance disparities, with models typically achieving higher accuracy for majority populations represented in the training data. In this study, we evaluated whether supervised domain adaptation methods—including balanced weighting, two-stage TrAdaBoost, feature augmentation with SrcOnly prediction, and linear interpolation—can mitigate these biases. Using the ABCD dataset, we assessed whether models trained on 80 different MRI measures from White American participants could generalize more effectively to African American participants. All domain adaptation methods reduced prediction error for African American participants, particularly for MRI modalities with large baseline disparities (e.g., structural MRI), while offering limited improvements where initial gaps were smaller (e.g., functional connectivity). Among the approaches, balanced weighting performed best and remained stable and beneficial even when only 10 African American participants were used to adapt the original model trained exclusively on White American participants. These findings suggest that simple, low-cost strategies can effectively reduce cross-ethnic performance gaps and improve equity in predictive neuroimaging, offering a practical path forward for future neuroimaging predictive biomarkers.

**Significant Statement:** Large-scale neuroimaging datasets increasingly enable machine-learning models to predict cognitive and clinical outcomes; however, these datasets are often ethnically/racially imbalanced. As a result, predictive models tend to generalize poorly to underrepresented populations. We demonstrate that, across 80 MRI phenotypes, a class of machine-learning approaches collectively known as supervised domain adaptation can substantially reduce cross-ethnicity disparities in neuroimaging-based cognitive prediction, even when only limited data from underrepresented groups are available. Among the methods evaluated, balanced weighting achieved the best performance while maintaining low computational cost. Together, these findings provide a practical and scalable framework for improving fairness and generalizability in neuroimaging-based machine learning under realistic conditions of ethnic/racial imbalance.

## Introduction

Recent progress in machine learning, combined with increasing access to large-scale neuroimaging datasets, has facilitated a growing number of applications using neuroimaging-based models to predict cognitive and clinical outcomes (Alhuwaydi, 2024; Finn et al., 2015; Sripada, Angstadt, et al., 2020; Sripada, Rutherford, et al., 2020; Sui et al., 2020; Tetereva et al., 2022, 2025; Vieira et al., 2022). Yet, most large-scale datasets consist of White participants from Europe and North America, raising concerns about fairness and generalisability to different ethnicities. Studies have documented performance disparities across ethnic/racial groups when machine-learning models are trained on imbalanced samples, demonstrating that training data imbalances and underlying population structure can lead to systematically poorer performance for underrepresented groups (Carey et al., 2024; Cross et al., 2024; Hasanzadeh et al., 2025; Ueda et al., 2024).

In the context of brain–behaviour modelling, such disparities are particularly concerning, as neuroimaging data are high-dimensional, noisy, and costly to acquire, and bias may reinforce existing inequities in neuroscience research and downstream applications. Prior work in large-scale neuroimaging cohorts has demonstrated that brain MRI-based models showed systematic differences in prediction error across ethnic groups when predicting cognitive-related outcomes (Adkins & Hanson, 2025; Li et al., 2022). These biases were not uniform across neuroimaging phenotypes. For instance, structural MRI showed larger cross-ethnic performance gaps than functional MRI (Lal Khakpoor et al., 2025).

Bias in predictive modelling can arise at multiple stages, spanning data collection, pre-processing, feature construction, model development, and post-deployment evaluation (Hasanzadeh et al., 2025). In principle, early-stage interventions— such as more diverse data collection or unbiased pre-processing pipelines — are likely to provide the most fundamental solutions. In practice, however, these approaches are often slow to implement, resource-intensive, or unavailable for large existing datasets. For instance, datasets from countries with more limited research budgets are likely to be smaller. As a result, there is a need to understand what can be achieved using currently available imbalanced data and standard pre-processing pipelines at later stages of the pipeline.

In our prior work (Lal Khakpoor et al., 2025), we addressed representation bias through prior-driven balanced and oversampling strategies. We showed that balancing sample size, by subsampling majority-group participants to match the size of minority groups, narrowed the gap in performance but left disparities significant among MRI phenotypes with greater bias, such as structural MRI . This suggests that the issue is not just "how many" samples, but rather differences in the feature**-**space relationship between ethnicity and cognition. Therefore, ensuring balanced samples in neuroimaging datasets improves performance but does not guarantee fairness across groups. Moreover, subsampling reduces statistical power, increases variance, and may compromise model stability (Rajput et al., 2023; Susan & Kumar, 2021)—an especially critical issue for neuroimaging data, where reliable estimation already requires large samples (Rajput et al., 2023). Balancing is also often impractical or infeasible when minority-group sample sizes are very small, a frequent scenario in real-world applications. This motivates the exploration of alternative methodological frameworks that can operate effectively under extreme sample imbalance and explicitly account for cross-group differences in data distributions.

One such framework is domain adaptation (DA). Domain adaptation methods are designed to improve model performance when training (source) and test (target) data come from different distributions (Farahani et al., 2021). In neuroimaging, DA techniques have been widely applied to address site effects, scanner differences, and protocol variability in multi-site studies as well as cross-task and cross-dataset classifications (Bayer et al., 2022; Cheng et al., 2015; Kushol et al., 2024; Liu et al., 2019; Panda et al., 2022; Wang et al., 2022; Zhang et al., 2020). These methods aim to reduce performance degradation when models trained on one domain are applied to another, without requiring full retraining on large target datasets. However, despite their extensive use in cross-site harmonization and generalization problems, domain adaptation approaches have not been examined as tools for mitigating ethnicity-related prediction gap in neuroimaging-based predictive models, particularly for regression-based prediction of continuous outcomes such as cognitive or behavioural measures.

Within a domain adaptation framework, ethnic groups can be conceptualized as distinct domains characterized by partially overlapping but non-identical distributions (Mo & Siepel, 2023; Zhou et al., 2021). In this view, the majority group can be treated as the source domain, while the minority group participants—constitutes the target domain. The primary objective here is not to equalize performance across ethnic groups, but to improve predictive accuracy for the target domain (i.e., the minority participants) using information learned from the source (i.e., the majority participants). This can be achieved by using a small number of labelled target samples — meaning target-domain examples that include the observed output (observed cognitive functioning scores, in the case of using MRI to predict cognitive functioning). These labelled examples from the target domain help “nudge” the source-trained model toward understanding the target data distribution (Guan & Liu, 2022). An effective adapted model should therefore be able to encode what generally works and how it differs for the target domain.

Beyond performance gain for the target domain, domain adaptation methods can also provide insight into the mechanisms underlying the performance gap, as comparing feature importance between adapted and non-adapted models can help identify which features change most in their contribution to prediction when adapting to the target domain.

In this study, we addressed the ethnicity-related performance gap in neuroimaging-based cognitive prediction as a domain adaptation problem using the Adolescent Brain Cognitive Development Study (ABCD) dataset. Treating White American (WA) participants as the source domain and African American (AA) participants as the target domain, we evaluate whether domain adaptation can improve neuroimaging-based predictions of cognitive functioning for the underrepresented group when only limited labelled target data are available—a setting in which balanced sampling is infeasible or undesirable. Here, we analysed 80 unimodal neuroimaging phenotypes and focused on those producing the largest and smallest performance gaps, defined as the difference in prediction error when a model trained on only white samples is tested on White-American vs. African-American samples. We systematically evaluated supervised domain adaptation methods that incorporated increasing numbers of labelled target samples and compared their performance against non-adapted models trained with the same data. We utilised four methods applicable to the regression task, including balanced weighting, two-staged TrAdaBoost, feature augmentation with SrcOnly prediction and linear interpolation. Figure 1 shows the overall design of the study.

**Figure 1.**
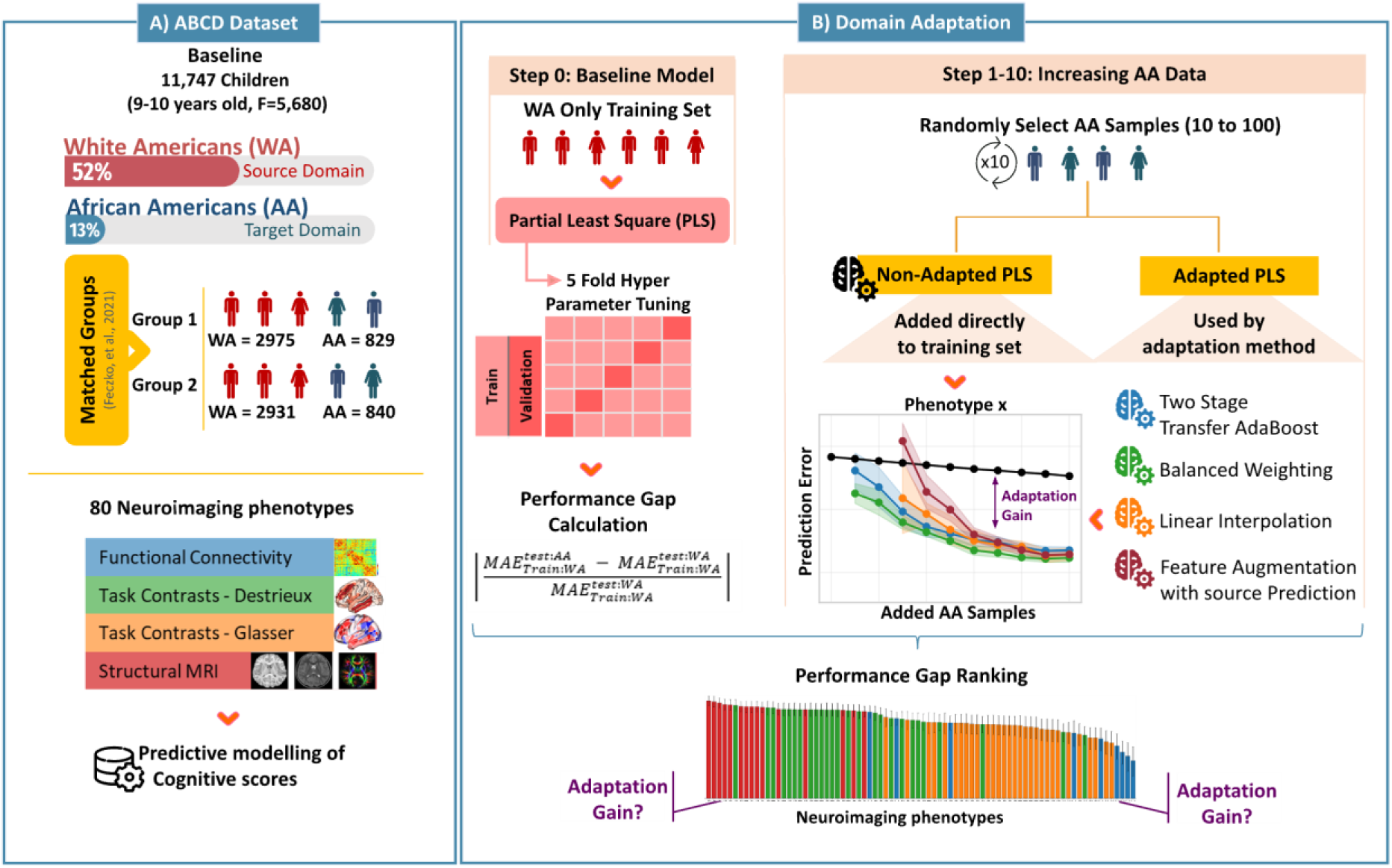
Overall Design. **(A)** Participants, matched groups and data extracted from the ABCD dataset. **(B)** Domain adaptation procedure.

By systematic evaluation of domain adaptation performance, this work aims to clarify whether and for which phenotypes domain adaptation can mitigate ethnicity-related performance gap in neuroimaging-based prediction of cognition. More broadly, it seeks to assess the practical utility of domain adaptation in neuroimaging-based predictive modelling with underrepresented populations.

## Methods

### Participants

Data were drawn from the Adolescent Brain Cognitive Development study (ABCD-5.1, DOI:10.15154/z563-zd24), a longitudinal neuroimaging project with a diverse U.S. cohort. The baseline cohort included 11,878 participants (48% female; 52% male) aged 9–10 years (Jernigan et al., 2018). As shown in Figure 1A, to investigate ethnicity-related generalisation, we focused on White-American and African-American participants drawn from the two largest matched groups within the ABCD-BIDS Community Collection (ABCC-2.0) (ABCD-BIDS Community). The ABCC provides matched groups based on clinical history (e.g., anaesthesia exposure), demographic variables (age, sex, race/ethnicity), and socioeconomic indicators (family income, household size, and family structure) (Feczko et al., 2021). For the purposes of the current study, we used the terms “race” and “ethnicity” interchangeably to refer to the characterisation of WA and AA groups.

### Cognitive measures

Cognitive performance was indexed using the NIH Toolbox total cognition composite score provided in the ABCD dataset (Akshoomoff et al., 2013; Jernigan et al., 2024) derived from seven tasks: Picture Vocabulary (Gershon et al., 2013), Flanker Inhibitory Control & Attention (Zelazo et al., 2013), List Sorting Working Memory (Tulsky et al., 2013), Dimensional Change Card Sort (Zelazo et al., 2013), Pattern Comparison Processing Speed (Carlozzi et al., 2013), Picture Sequence Memory (Bauer et al., 2013), and Oral Reading Recognition (Gershon et al., 2013). Together, these tasks provide a comprehensive assessment of cognitive function across multiple domains.

### Neuroimaging Data and Processing

#### Data Sources

Neuroimaging phenotypes were derived from the Adolescent Brain Cognitive Development (ABCD) Study and integrated across two complementary sources: (i) post-processed tabulated summary measures released by the ABCD consortium (version 5.1) (Jernigan et al., 2024) and (ii) minimally pre-processed data from the ABCD-BIDS Community Collection (ABCC-2.0) formatted according to the Brain Imaging Data Structure (BIDS) standard (ABCD-BIDS Community).

The ABCD tabulated datasets provide quality-controlled regional measures and task contrasts parcellated using the Destrieux cortical atlas (Destrieux et al., 2010) and FreeSurfer subcortical segmentation (Fischl et al., 2002), as described in Hagler et al. (2019) and Jernigan et al. (2024).

To extend spatial resolution and broaden structural and connectivity coverage beyond these summaries, we conducted additional processing on ABCC data using outputs from fMRIPrep (v20.2.0) and the Human Connectome Project (HCP) minimal pre-processing pipeline. The ABCC includes pre-processed MRI images, confounds tables generated by fMRIPrep, and CIFTI-formatted surface data from the HCP pipeline.

In total, this pipeline yielded 80 distinct neuroimaging phenotypes (Supplementary Figure 1), spanning task contrasts, functional connectivity, structural MRI, and diffusion imaging.

#### Quality Control and Exclusion Procedures

Quality control followed ABCD-recommended criteria (Jernigan et al., 2024). Exclusion flags were applied at the level of individual neuroimaging features based on image quality metrics, MR neurological screening, and task performance indicators (Hagler et al., 2019).

We did not perform listwise deletion. Instead, exclusions were applied at the level of the neuroimaging phenotype. For example, if a participant failed quality criteria for the N-back task fMRI but met criteria for other modalities, only the N-back phenotype was excluded.

We excluded Site 22 due to limited sample size and corrected site ID errors (Garavan et al., 2018). Additionally, 58 participants with reported vision problems were excluded (Luciana et al., 2018). Final sample sizes ranged from approximately 4,000 to 11,177 participants, depending on modality and phenotype.

### Neuroimaging Phenotypes

#### Functional MRI (fMRI) - Task-Evoked Activation (56 Sets)

Task-evoked BOLD responses were derived from three ABCD tasks: Emotional N-back task (Cohen et al., 2016), Monetary Incentive Delay (MID) task (Knutson et al., 2000), and Stop Signal Task (SST) (Logan, 1994). We used 26 precomputed contrasts from the ABCD tabulated releases (9 N-back, 10 MID, 7 SST), parcellated using the Destrieux cortical atlas (Destrieux et al., 2010) and FreeSurfer subcortical segmentation (Fischl et al., 2002; Hagler et al., 2019). We computed an additional 30 contrasts (10 per task) using ABCC data and a first-level General Linear Model (GLM) implemented primarily in Nilearn (Abraham et al., 2014; Nilearn contributors et al., 2026) and Nibabel (Brett et al., 2024).

#### Event Processing

Task event files in E-Prime format were converted to BIDS-compatible TSV files using the eprimetotsv.py script (Demidenko, 2024). Event onsets were aligned relative to post-calibration task start times.

#### Pre-processing and Nuisance Regression

Nuisance regressors were selected based on prior recommendations (Ciric et al., 2017; Gotts et al., 2013). Included confounds were Cosine drift regressors to remove low-frequency noise (Ciric et al., 2017), ten anatomical CompCor components derived from white matter and CSF masks (Muschelli et al., 2014), and six rigid-body motion parameters (translations and rotations) and their first derivatives (Maknojia et al., 2019). Specific motion regressors included rot_x, rot_y, rot_z, trans_x, trans_y, trans_z and their first derivative terms. Non-steady-state volumes were removed prior to modeling. The number of removed TRs depended on scanner type: Siemens and Philips (8 TRs), GE DV25 (5 TRs), GE DV26 (16 TRs) (Jernigan et al., 2024).

#### General Linear Model Estimation

First-level design matrices were constructed using the SPM canonical hemodynamic response function. Time series were standardized, and General Linear Models (GLMs_ were estimated using ordinary least squares (OLS). Contrast matrices were defined to isolate task-specific cognitive processes. Run-level beta coefficients were estimated and combined across runs using variance-weighted fixed effects. Final contrast maps were parcellated into 379 regions (360 Glasser cortical parcels (Glasser et al., 2016) and 19 FreeSurfer subcortical regions) to yield regional effect sizes.

### Functional Connectivity (9 Sets)

Functional connectivity (FC) phenotypes were derived from both ABCD tabulated releases and ABCC processing. From ABCD Tabulated Resting-State FC data, we included: 1) Temporal variance (amplitude of low-frequency fluctuations) across 333 Gordon cortical parcels (Gordon et al., 2016) and 19 subcortical regions. 2) Subcortical-to-network connectivity (average correlations between 19 subcortical regions and 13 cortical networks). 3) Within- and between-network cortical connectivity across 13 large-scale networks (Jernigan et al., 2024). Using ABCC data, we computed six 379-region FC matrices for Rest, N-bac, MID, SST, Multitask FC (concatenation across tasks) (Cole et al., 2021), and General FC (rest + all tasks concatenated) (Elliott et al., 2019). Concatenation-based FC measures were motivated by evidence that aggregating across tasks improves reliability and behavioural prediction (Cole et al., 2014, 2019; Elliott et al., 2019).

### Motion and Outlier Handling

High-motion TRs were identified using Framewise displacement (FD > 0.5) (Power et al., 2013) and DVARS > 1.5 (Afyouni & Nichols, 2018; Phạm et al., 2023). Flagged TRs were included as nuisance regressors (Power et al., 2014). Participants were excluded if more than 50% of timepoints were flagged. A relatively lenient FD threshold was adopted to balance artifact mitigation with signal preservation in this developmental, multiband dataset (Addeh et al., 2023; Phạm et al., 2023; Power et al., 2019).

### Connectivity Estimation

Nuisance regressors were selected and included in the design matrix, similar to task contrasts. For task FCs, the events were also included in the design matrix. After nuisance regression, residual time series were extracted, de-spiked (values >3 SD replaced with mean; (Costabel & Müller-Petke, 2014), concatenated across runs, and parcellated into 379 regions. Functional connectivity was computed using Pearson correlation, followed by Fisher z-transformation and vectorization.

#### Structural MRI (sMRI) and Diffusion MRI (16 Sets)

Structural phenotypes were derived primarily from FreeSurfer processing of T1- and T2-weighted images (Fischl et al., 2002). Cortical regions were parcellated using the Destrieux atlas, and subcortical regions using FreeSurfer segmentation. We included 15 structural phenotypes that captured complementary aspects of morphology and tissue properties. FreeSurfer summations comprised five global metrics: estimated intracranial volume, total cortical gray matter volume, total cortical white matter volume, total subcortical gray matter volume, and brain segmentation volume-to-intracranial volume ratio (Tetereva et al., 2022). T1 and T2 summary phenotypes capture broader regional and global signal characteristics sensitive to tissue composition, including myelination and water content (Jørgensen et al., 2016; Xu et al., 2024), complementing localized surface-based intensity measures.

Diffusion phenotypes consisted of fractional anisotropy (FA) extracted from 23 major white-matter tracts identified using AtlasTrack (Hagler Jr et al., 2009). FA quantifies directional water diffusion and indexes white-matter microstructural organization (Alexander et al., 2007) (Alexander et al., 2007). The processing steps are explained in Hagler and colleagues (2019).

See Supplementary Figure 1 for the list of phenotypes and their corresponding sample sizes.

### Ethnicity-related Performance Gap

To quantify cross-ethnicity misprediction for each neuroimaging phenotype, we computed a performance gap based on differences in prediction error when models trained on one ethnicity were evaluated on the same-ethnicity versus different-ethnicity test samples. Specifically, this metric captures the relative increase in prediction error when a model trained on White Americans is applied to African Americans, compared to its performance on White Americans test data (higher values indicate more misprediction for AA). This measure provides an index of cross-ethnicity generalisation error and was used to stratify phenotypes by baseline bias, enabling evaluation of domain adaptation under varying levels of misprediction. The performance gap was defined as:

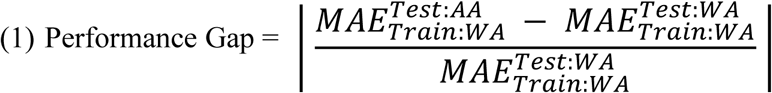

Superscript shows the test group that MAE is calculated for, and subscript indicates the group that the model is trained on.

### Domain Adaptation

We evaluated four machine learning-based supervised domain adaptation methods implemented in the ADAPT Python library (de Mathelin et al., 2021), all of which are applicable to regression settings and allow explicit incorporation of labelled target-domain samples. Unlike unsupervised methods that only align feature distributions, these supervised approaches utilize labelled target data to address shifts in outcome (cognitive scores) distributions *p(y)*, marginal shifts in feature space (input MRI data) *p(x)*, and feature-outcome conditional shifts *p*(*y*|*x*) either directly or indirectly (Farahani et al., 2021; Katz et al., 2024). To ensure computational feasibility and consistency across analyses, adaptation hyperparameters were fixed across steps and phenotypes. These values were selected based on an initial exploratory grid search on validation performance. We selected the most frequently optimal parameter setting and applied it uniformly throughout the study.

### Balanced Weighting

Balanced Weighting adjusts the importance of source and target samples during training by assigning higher weights to target-domain instances relative to source-domain instances. By increasing the influence of target samples, Balanced Weighting encourages the learned model to better reflect the target-domain distribution without discarding source-domain data (Wu & Dietterich, 2004). In this study, Balanced Weighting allows examination of whether reweighting a small number of AA samples can improve prediction relative to naïve inclusion in non-adapted training. The ratio parameter, which corresponds to the importance given to the target labelled data, was set to 0.8. This was fixed across adaptation steps and was decided based on an exploratory grid search.

### Two-stage TrAdaBoost

a sophisticated instance-based boosting algorithm that iteratively reweights samples. It identifies and down-weights misleading source samples—those whose brain-cognition associations contradict the patterns observed in the target AA domain—while boosting the influence of informative target samples (Pardoe & Stone, 2010). The number of boosting iterations for the first and second stages was fixed at 10 and 20, respectively, with a learning rate of 0.1.

### Feature Augmentation with SrcOnly Prediction (PRED)

A feature-based adaptation method where a model is first pretrained on the source domain. The resulting source-only predictions of target labelled data are then used as an augmented feature to train a second-stage model on the target data. This approach allows the model to leverage source-group knowledge as an informative prior while specifically learning target-group residuals (Daumé, 2009).

### Linear Interpolation

A parameter-based adaptation strategy that creates a new model by interpolating between the parameters of a source-only model and a target-only model. By weighting contributions from both domains, this method seeks an optimal parameter set that preserves generalizable source knowledge while accounting for target-specific signals. This approach does not explicitly reweight individual samples but instead interpolates between source and target solutions as the proportion of labelled target data increases (Daumé, 2009). The proportion of target-domain data used to train the target-only model was set to 0.7. The remaining target data were used to estimate the interpolation parameter.

### Baseline Estimator

Partial Least Squares (PLS) was used as the estimator for adaptation methods and as the standard non-adapted model. PLS offers several advantages that are particularly relevant for the present study. First, it is well suited to high-dimensional, highly collinear neuroimaging data, such as functional connectivity features, where the number of predictors greatly exceeds the number of observations (Krishnan et al., 2011). Second, PLS provides a low-dimensional latent representation that is directly optimized for predicting the outcome (Krishnan et al., 2011), which facilitates interpretability of feature reweighting across domains. Third, PLS is computationally efficient, given the large number of phenotypes, adaptation methods, and resampling steps evaluated in this study. To optimize the number of PLS components, we performed a grid search with five-fold cross-validation, capping the number of components at 30. The optimal model was selected based on the negative mean squared error (NMSE).

### Experimental design

The experimental design followed a structured, stepwise adaptation framework to evaluate how model performance on AA participants changes as increasing amounts of labelled target data are incorporated (Figure 1B).

In Step 0, a baseline PLS regression model was trained using only WA participants from the training set (i.e., either matched group 1 or 2). This model represents the source-only scenario and serves as the reference point for all subsequent comparisons.

In Steps 1–10, labelled AA samples were incrementally added to the training process. At each step, a fixed number of AA participants (ranging from 10 to 100 in increments of 10) were randomly selected from the training pool (i.e. either matched group one or two). These samples were either (i) included directly in the training of a non-adapted PLS model or (ii) provided as labelled target-domain data to the domain adaptation methods. This design allows direct comparison between naïve data inclusion and adaptation-based modelling under matched conditions.

For each step, the random selection of AA samples was repeated 10 times to account for sampling variability. Model performance was evaluated on a held-out AA test set that was not used during training or adaptation (e.g., from matched group 2 when the model was trained on matched group 1). Predicted values, error metrics, and model coefficients were saved for each repetition and step for subsequent analysis.

### Evaluation metrics

For each adaptation step, MAE values were obtained across repeated random selections of labelled AA samples. To compare adapted models with non-adapted training at each step, paired t-tests were conducted across matched repetitions, comparing MAE values from adapted and non-adapted models trained on the same randomly selected AA samples.

For Linear Interpolation and PRED methods, statistical comparisons were restricted to adaptation steps with N = 30–100, as these approaches involved training target-only models and were unstable at smaller target sample sizes. In contrast, Balanced Weighting and Two-stage TrAdaBoost were stable from N = 10.

To summarize adaptation benefits across target sample sizes, we computed the area under the improvement curve (AUIC), defined as the integral of ΔMAE_(N)_ = MAE_non-adapted(N)_ − MAE_adapted(N)_ across adaptation steps starting from N=30. Starting at N=30 allowed us to compare AUIC across all four adaptation methods. This scalar measure provides a compact summary of cumulative adaptation benefit across increasing availability of labelled AA data, without assuming independence between steps. The same cumulative adaptation benefit was also calculated using R-squared values. We computed Pearson’s correlation between performance gap scores and cumulative adaptation AUIC gains (averaged across adaptation methods) to test whether adaptation benefit scales with performance gap severity. This analysis evaluates whether adaptation acts primarily as a gap-mitigation mechanism rather than providing uniform gains across phenotypes.

To compare the effectiveness of the four domain adaptation approaches in reducing the performance gap, cumulative AUIC gains were compared across phenotypes using repeated-measures nonparametric analyses. An omnibus Friedman test was first performed to assess overall differences between methods. Pairwise post-hoc comparisons were then conducted using two-sided Wilcoxon signed-rank tests with Holm correction for multiple comparisons. Effect sizes for pairwise comparisons were quantified using paired Cohen’s *d*.

### Model interpretation

To examine how adaptation alters outcome-relevant feature weighting, we tracked changes in PLS regression coefficients across adaptation steps for the adaptation method with the largest gain value. For each MRI phenotype, we computed coefficient differences between models trained with no AA data (Step 0) and models trained with 100 labelled AA samples (step 10).

To isolate adaptation-specific effects, we employed a difference-of-differences formulation: (i) the change in coefficients for the non-adapted PLS model after adding 100 AA samples was computed, (ii) the corresponding change for an adapted model was computed, and (iii) the difference between these two changes was taken as an index of adaptation-related reweighting beyond naïve data inclusion. Higher values reflect larger adaptation-specific shifts in the feature’s signed contribution to the predicted outcome (PLS coefficient), beyond changes induced by adding AA data alone.

Coefficient changes were visualized by mapping features back onto brain space for each phenotype. These analyses are descriptive and intended to characterize patterns of feature reweighting associated with adaptation; they do not support causal inference regarding the sources of bias.

## Results

### Performance Gap Ranking

The ranking of phenotypes based on the gap metric revealed systematic differences across modalities, as shown in Figure 2. Functional connectivity measures exhibited the smallest gap values, indicating that models trained exclusively on WA participants showed the least degradation in performance when applied to AA participants relative to WA test performance.

**Figure 2.**
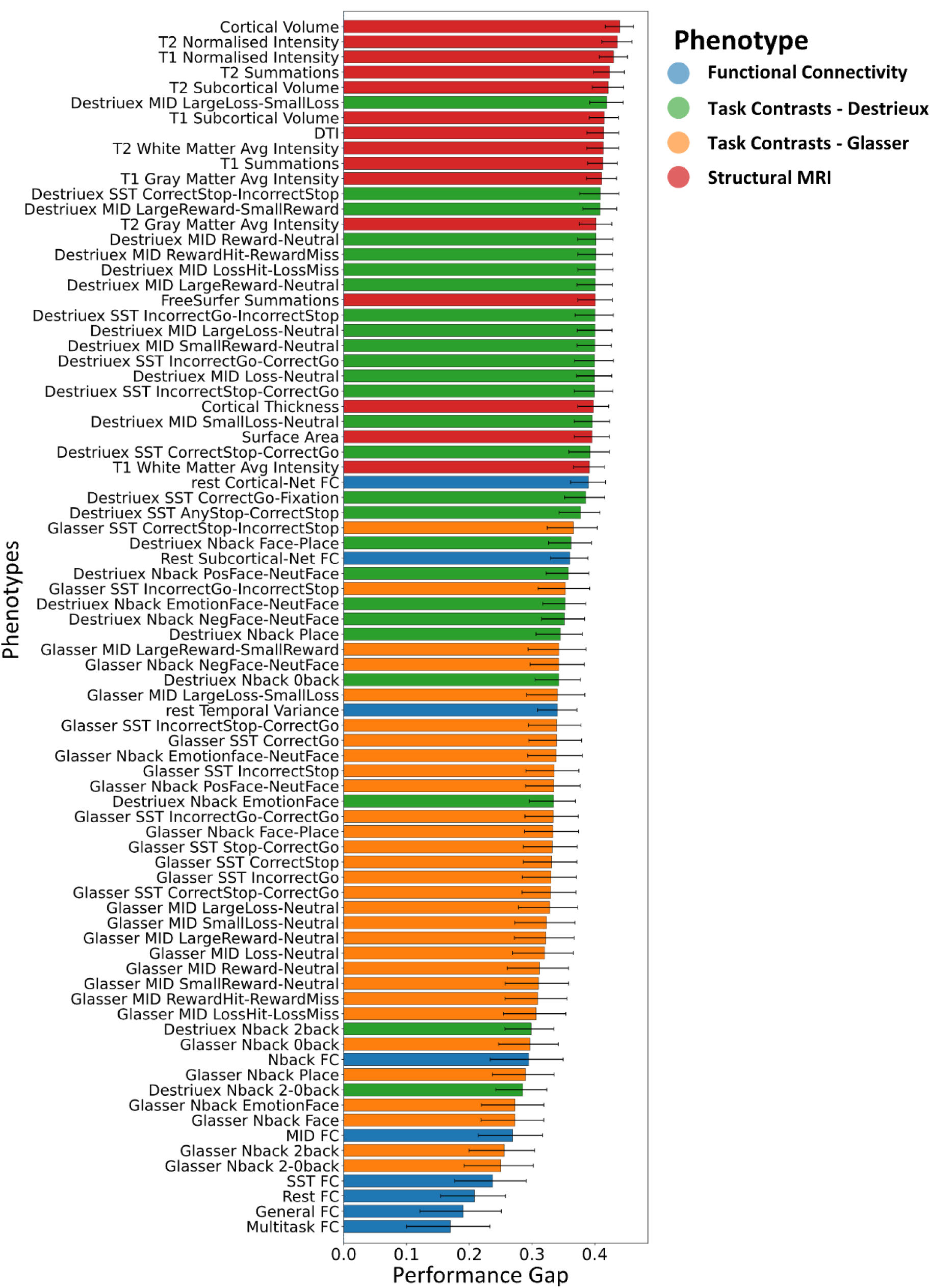
The gap in predictive performance (MAE) of the Baseline Model (trained only on WA) when tested on WA versus AA participants in the test set. Confidence intervals were estimated using non-parametric bootstrap resampling (5,000 iterations) at the subject level, with resampling performed independently within AA and WA groups. For each bootstrap sample, MAEs and the gap metric were recomputed, and 95% confidence intervals were derived using the percentile method.

This pattern was followed by task-based fMRI contrast measures derived from the Glasser parcellation, particularly Nback task contrasts, which also demonstrated relatively low gap values. Across task-based contrasts more broadly, most phenotypes occupied an intermediate range of the ranking, with Glasser-parcellated phenotypes exhibiting lower gap values compared to Destrieux-based phenotypes.

In contrast, sMRI and DTI phenotypes consistently showed the largest gap values, indicating greater disparity in model performance between AA and WA participants.

### Domain adaptation performance

We first evaluated whether supervised domain adaptation improves prediction performance for AA participants relative to non-adapted PLS models trained with naïve inclusion of AA samples.

To summarize adaptation effects across target sample sizes, we computed the area under the improvement curve (AUIC) for each of 80 MRI phenotypes and adaptation methods. Figure 3a illustrates adaptation gains of each method across phenotypes ranked by their performance gap score. AUIC values revealed a clear structure across phenotypes: phenotypes with a larger gap showed larger cumulative gains, whereas those with a smaller gap showed smaller improvements. Connectivity phenotypes consistently showed near-zero or negative AUIC values, indicating minimal or absent cumulative benefit from adaptation. A similar pattern was observed when we calculated the adaptation benefit based on changes in the coefficient of determination (R2), as shown in Supplementary Figure 2. The performance gap score was significantly correlated with average gain from adaptation (*r=*0.79, *p<*0.001) (Figure 4b).

**Figure 3.**
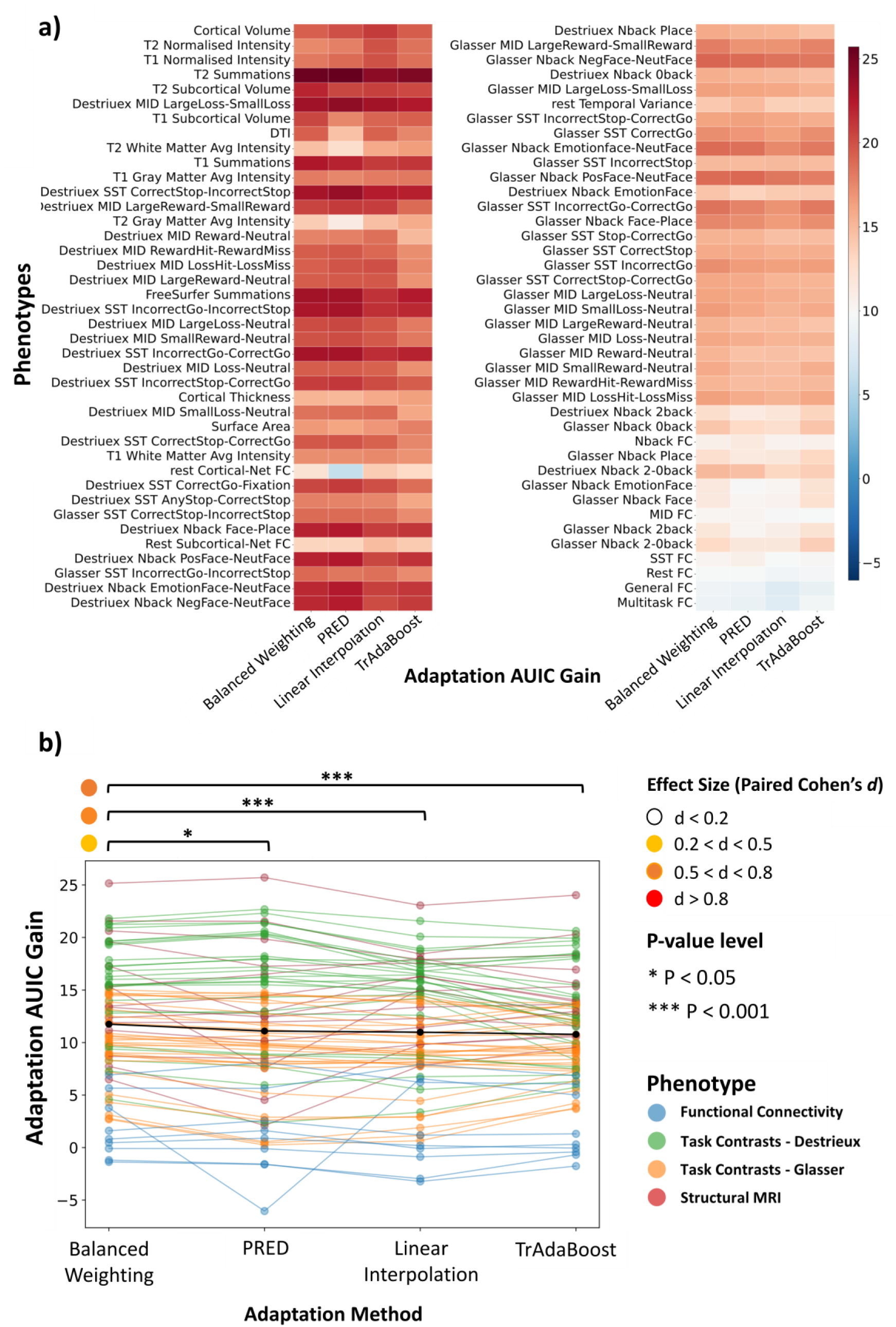
Adaptation Benefits. **(a)** The cumulative AUIC gains of each adaptation method across all phenotypes, sorted by their performance gap scores, starting from phenotypes with the largest gap at the top left to phenotypes with the smallest gap at the bottom right. A positive AUIC means the adapted model produced a cumulative reduction in MAE as AA labels increased. **(b)** Repeated-measures comparison of adaptation-method AUIC gain across neuroimaging phenotypes. Each line represents an individual phenotype tracked across adaptation methods, illustrating within-phenotype changes in performance gain. The black line shows the average gain across phenotypes for each method. Colored points indicate phenotype-specific AUIC values for each method. Significance levels from Pairwise Wilcoxon signed-rank tests with Holm correction and paired Cohen’s *d* are shown.

**Figure 4.**
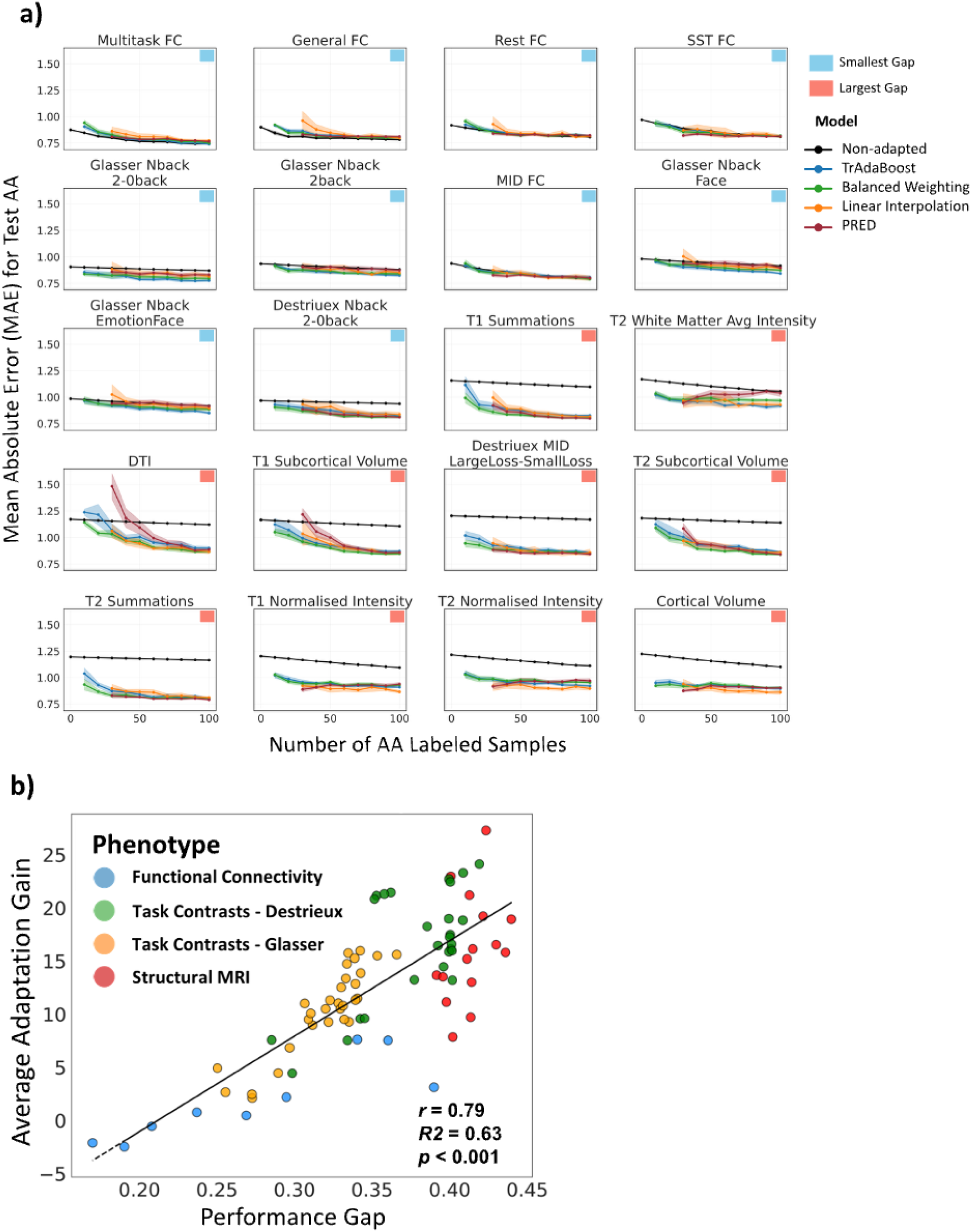
**(a)** Changes in AA test prediction error as increasing numbers of labelled AA participants were incorporated into model training, either by direct inclusion (non-adapted) or via adaptation. Points show the mean MAE across repeated random AA subsamples at each target sample size, and shaded bands indicate approximate 95% confidence intervals across repetitions (±1.96 SD/n). Results are shown for the ten phenotypes with the largest and smallest gap scores. **(b)** The scatter plot of the correlation between average cumulative AUIC gain and performance gap across phenotypes.

Across MRI phenotypes with a larger gap, all adaptation methods showed significant reductions in MAE relative to non-adapted training, as shown in Figure 4a, and these reductions persisted as additional target samples were incorporated (*p* < 0.05). Across MRI phenotypes with a lower gap, task-based contrasts also showed improvements with adaptation; however, the magnitude of MAE reduction was smaller, and adapted and non-adapted performance lines were closer throughout the adaptation range. Nevertheless, adapted models often achieved modest but consistent reductions in MAE. In contrast, connectivity phenotypes showed little to no benefit from adaptation, where performance curves for adapted and non-adapted models largely overlapped across adaptation steps, and in some cases, adapted models exhibited slightly higher MAE before converging to similar values at larger sample sizes. The change in prediction errors across adaptation steps for each of 80 phenotypes is depicted in Supplementary Figure 3.

### Comparison across domain adaptation methods

We used a Friedman test to determine whether the largest cumulative reduction in MAE, as quantified by the Area Under the Improvement Curve (AUIC), differed significantly across Adaptation methods. The results showed a significant main effect of Adaptation methods, *χ*^2^(3) = 40.85, *p* = 7.05×10−9 (Figure 3b). Post hoc comparisons using the Wilcoxon signed-rank tests with Holm correction revealed that Balanced Weighting (*M* = 11.75, *SD* = 6.04) lead to significantly higher AUIC than PRED (*M* = 11.10, *SD* = 6.76, corrected *p* = .02, paired Cohen’s *d* = .36), Linear Interpolation (*M* = 10.99, *SD* = 5.99 corrected *p* < .001, paired Cohen’s *d* = .54), and Two-stage TrAdaBoost (*M* = 10.78, *SD* = 5.36 corrected *p* < .001, paired Cohen’s *d* =.61). However, there was no statistically significant difference in AUIC between PRED and Linear Interpolation (corrected *p* = .07, paired Cohen’s *d* = .05) and between PRED and Two-stage TrAdaBoost (corrected *p* = .07, paired Cohen’s *d* = .12) and between Linear Interpolation and and Two-stage TrAdaBoost (corrected *p* = .74, paired Cohen’s *d* =.11).

It is important to note that when the number of target samples was fewer than 30, PRED and Linear Interpolation were unstable and could not be reliably trained. In contrast, under the constant-strategy setting, Balanced Weighting and Two-stage TrAdaBoost remained stable and could be applied with as few as 10 target samples. Therefore, AUIC was computed starting from 30 target samples onward.

### Adaptation-related changes in PLS feature weights

Finally, we examined how domain adaptation altered outcome-relevant feature weighting by comparing changes in PLS coefficients between Step 0 and Step 100 for Balanced Weighting as the method with the highest average gain. Across phenotypes, the magnitude of adaptation-related coefficient changes varied substantially, as indicated by the range of color bars in Figure 5. Phenotypes with the largest gap exhibited the largest coefficient differences, while task-based phenotypes with smaller gap scores showed smaller changes, and connectivity phenotypes exhibited minimal coefficient differences, with maximum values below 0.001.

**Figure 5.**
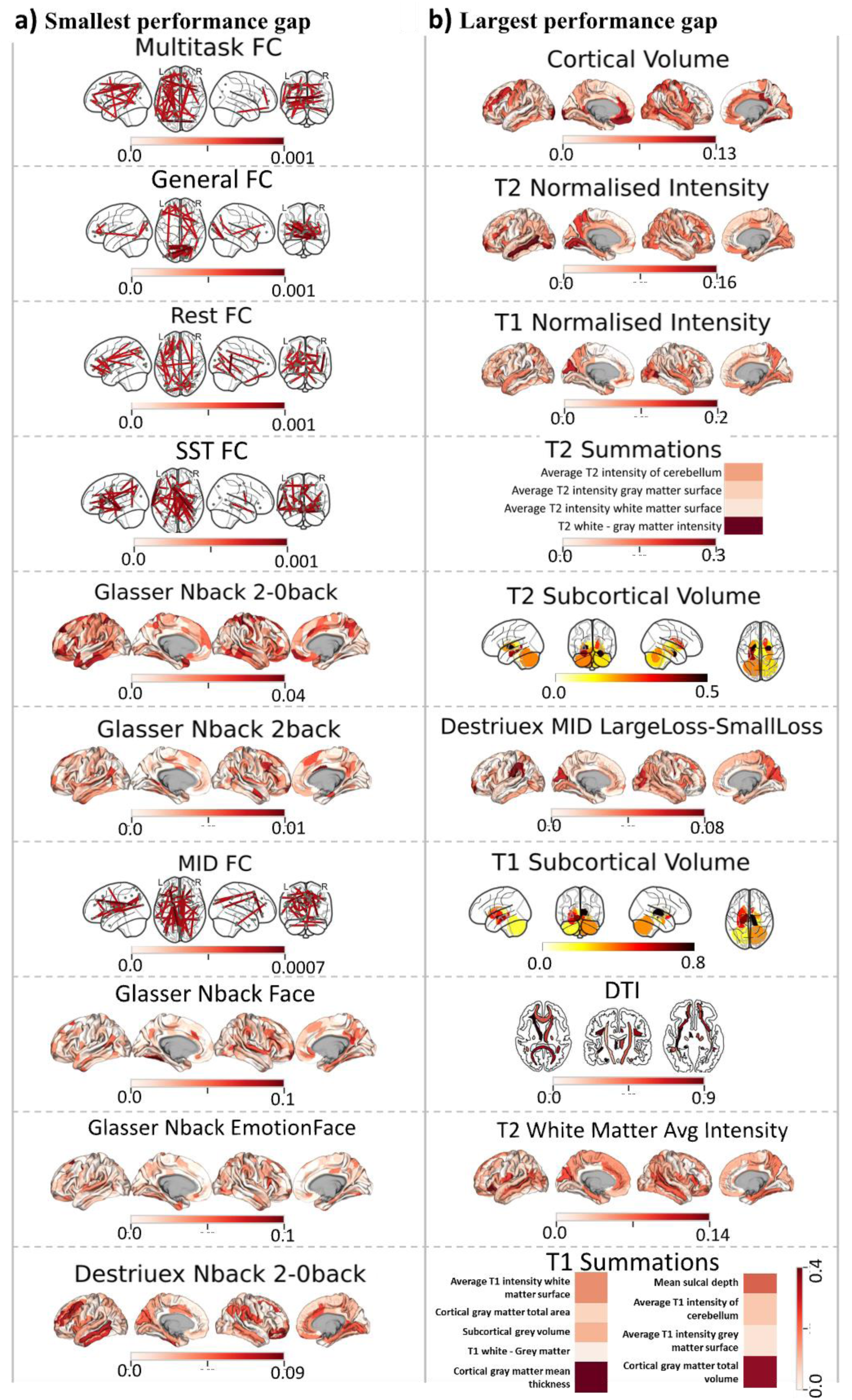
Adaptation-specific feature re-weighting across top ten phenotypes with the **(a)** smallest and **(b)** largest performance gap. This was estimated via a difference-of-differences framework comparing coefficient changes in adapted (Balanced Weighting) and non-adapted PLS models after adding 100 AA samples relative to the baseline (no AA added).

## Discussion

This study represents one of the first efforts to apply supervised domain adaptation to regression-based neuroimaging prediction to reduce ethnicity bias in neuroimaging-based cognitive prediction. We examined whether these methods could reduce ethnicity-related performance disparities in cognitive prediction, and identified the conditions under which they are most effective. Our systematic comparison of domain-adaptation strategies demonstrates that adaptation can improve prediction accuracy for AA participants even with limited labelled data, although the magnitude of improvement varies across phenotypes. Among the four strategies evaluated, balanced weighting achieved the best overall performance while maintaining low computational cost, highlighting its promise as a practical approach for reducing ethnicity bias in neuroimaging-based cognitive prediction.

### Phenotypes differ in performance gap across ethnicities

The ranking of performance gap scores indicates that functional connectivity phenotypes are comparatively more robust to cross-ethnicity generalisation when models are trained on WA participants. In contrast, structural and diffusion-derived features show greater sensitivity to cross-group differences, exhibiting larger increases in prediction error under cross-ethnicity evaluation.

### Supervised domain adaptation improves AA prediction with limited labelled data

Across most MRI phenotypes—particularly those with the largest gap —supervised domain adaptation led to significant reductions in prediction error on held-out AA test samples compared with naïve inclusion of the same labelled AA data in non-adapted PLS models. Notably, for Balanced Weighting and Two-stage TrAdaBoost, improvements were evident with as few as 10 labelled AA samples and increased as more target data became available.

At the same time, improvement trajectories varied substantially across MRI phenotypes. Some showed steep early declines in error, while others exhibited gradual or negligible change. This heterogeneity indicates that adaptation effectiveness cannot be inferred from representing sample size alone and instead reflects phenotype-specific properties of the underlying data-generating process.

The strongest overall improvements were observed for the instance-based adaptation approach of Balanced Weighting. This method leverages a small amount of labelled target-domain data to reweight source samples during training, allowing the model to place greater emphasis on source instances that are more representative of the target population. Despite its simplicity, Balanced Weighting achieved the largest overall reduction in performance disparities across phenotypes. Unlike Two-stage TrAdaBoost, which iteratively updates sample weights over multiple boosting rounds, Balanced Weighting applies a direct reweighting strategy, making it computationally efficient and straightforward to implement. Although the absolute performance differences between adaptation methods were modest, Balanced Weighting overall outperformed the other approaches and required substantially less computational overhead than Two-stage TrAdaBoost. In contrast, methods relying primarily on source-informed target predictions or parameter interpolation (PRED and Linear Interpolation) were not applicable in very small-sample settings, requiring more labelled target samples (at least 30) before adaptation could be performed. However, once sufficient target data were available, these approaches achieved performance improvements comparable to those of the instance-based methods.

### Adaptation reweights feature contributions

Analysis of PLS coefficient changes further supports this interpretation. Phenotypes that benefited most from adaptation exhibited larger adaptation-specific changes in regression coefficients beyond those induced by naïve inclusion of AA data. In contrast, phenotypes with little or no adaptation gain showed minimal additional changes in the coefficient.

This correspondence suggests that successful adaptation meaningfully alters the effective weighting of features toward the target domain when a source–target mismatch exists, resulting in both larger coefficient shifts and improved prediction. When the non-adapted model is already close to target-optimal, little reweighting occurs, and little improvement is observed. Adaptation-related changes in PLS weights were most consistently observed in orbitofrontal and cingulate cortices, with more distributed effects in visual and sensorimotor regions, based on cortical volume and normalized intensity measures. Subcortically (T1 and T2 subcortical volumes), the largest effects were concentrated in striatal structures, with additional involvement of thalamic and medial temporal regions (hippocampus and amygdala). These analyses are descriptive and do not imply causal involvement of specific brain regions or connections; rather, they characterize how adaptation redistributes predictive emphasis across features as a function of phenotype-specific mismatch.

### Practical implications

These findings have direct implications for fairness and generalisability in neuroimaging-based predictive modelling. They indicate that ethnicity-related performance gap is heterogeneous in origin and that mitigation strategies must be matched to the structure of domain differences. Supervised domain adaptation is most effective for phenotypes with ethnicity-specific predictive disparities. Although this study focused on ethnicity-related differences, the same framework holds promise for other underrepresented groups, including clinical populations, rare conditions, and sociodemographically stratified cohorts, particularly for continuous behavioural or cognitive outcomes for which model generalisability is rarely investigated.

In applied neuroimaging settings, obtaining balanced, exhaustive training data is often infeasible. Imaging protocols, scanners, and acquisition characteristics evolve over time; sensitive or clinical-labelled outcomes are costly and limited; and models trained on highly heterogeneous data may sacrifice accuracy by relying on overly coarse features to ensure robustness (Wang et al., 2022). Within these constraints, domain adaptation provides a data-efficient means of improving prediction for underrepresented groups by explicitly preserving transferable knowledge from the source domain while adjusting for target-specific shifts. Under scarce target-sample regimes, instance-based approaches, particularly, balanced weighting offer robust gains with minimal overhead. Moreover, our prior work has shown that structural features continue to favour WA populations even under balanced sampling (Lal Khakpoor et al., 2025). Given the widespread clinical availability of structural MRI, the ability of adaptation techniques to improve performance using small, labelled target samples is therefore particularly promising for real-world deployment.

Rather than framing fairness solely as equalizing performance across groups, these techniques enable the development of models optimized for peak performance within specific target groups, particularly when biological, environmental, or pre-processing factors inherently amplify domain distinctions in feature or outcome distributions that might not otherwise exist. This approach serves as a pragmatic solution in the absence of methods to fully eliminate such confounding factors.

### Limitations and future directions

Several limitations warrant consideration. First, we deliberately constructed a highly skewed adaptation scenario by restricting the number of labelled AA samples used for training to between 10 and 100, despite the availability of substantially larger AA samples in the full dataset. This design choice was intentional and motivated by our aim to evaluate the effectiveness of domain adaptation in settings where balanced sampling is infeasible or statistically undesirable. In real-world neuroimaging studies, minority groups are often severely underrepresented, and subsampling majority groups to achieve balance can lead to substantial loss of power and model instability. Our results, therefore, speak specifically to the utility of adaptation under extreme imbalance, rather than to the limits of performance achievable with fully balanced data.

Second, our analyses focused on unimodal neuroimaging phenotypes and linear regression models. This choice reflects both the high dimensionality of MRI data and the regression nature of the cognitive prediction task, and it enabled us to evaluate domain adaptation using well-established, widely validated supervised methods with strong theoretical and empirical grounding. However, this design limits direct generalization to multimodal or nonlinear modelling frameworks. More complex deep learning–based adaptation and representation learning approaches were therefore not explored and remain an important direction for future work (Liang et al., 2023; Stan & Rostami, 2023), with the present study providing a principled baseline for such extensions.

Finally, we examined a single composite cognitive score, derived from multiple cognitive tasks, within one large cohort. Extension of this framework to other behavioural, clinical, or developmental targets—particularly those with differing noise characteristics or domain sensitivities—will be important for assessing the utility of domain adaptation.

## Conclusion

In summary, this study demonstrates that supervised domain adaptation can substantially mitigate ethnicity-related performance disparities in regression-based neuroimaging prediction while requiring only a small amount of labelled target-population data.

Across diverse MRI phenotypes, adaptation-based models consistently produced larger improvements in predictive performance than simply adding the same target samples into conventional training pipelines. These improvements were achieved using computationally efficient methods—particularly balanced weighting—without requiring fully balanced multi-ethnic training datasets or more computationally intensive mitigation strategies, such as ethnicity-specific or ethnicity-neutral template construction, specialised pre-processing pipelines, or large-scale deep learning fine-tuning. Notably, the largest adaptation benefits were observed for MRI phenotypes showing the greatest baseline cross-ethnicity performance disparities, suggesting that supervised adaptation may be especially valuable for high-bias predictive features.

Overall, these findings highlight the value of targeted adaptation strategies for improving model performance in underrepresented groups without requiring fully balanced datasets. More broadly, they support the use of domain adaptation as a principled framework for enhancing fairness and generalisability in neuroimaging-based prediction of continuous behavioral outcomes.

## Acknowledgments

Data used in the preparation of this article were obtained from the Adolescent Brain Cognitive Development (ABCD) Study (https://abcdstudy.org), held in the NIMH Data Archive (NDA). This is a multisite, longitudinal study designed to recruit more than 10,000 children age 9-10 and follow them over 10 years into early adulthood. The ABCD Study® is supported by the National Institutes of Health and additional federal partners under award numbers U01DA041048, U01DA050989, U01DA051016, U01DA041022, U01DA051018, U01DA051037, U01DA050987, U01DA041174, U01DA041106, U01DA041117, U01DA041028, U01DA041134, U01DA050988, U01DA051039, U01DA041156, U01DA041025, U01DA041120, U01DA051038, U01DA041148, U01DA041093, U01DA041089, U24DA041123, U24DA041147. A full list of supporters is available at https://abcdstudy.org/federal-partners.html. A listing of participating sites and a complete listing of the study investigators can be found at https://abcdstudy.org/consortium_members/. ABCD consortium investigators designed and implemented the study and/or provided data but did not necessarily participate in the analysis or writing of this report. This manuscript reflects the views of the authors and may not reflect the opinions or views of the NIH or ABCD consortium investigators. The ABCD data repository grows and changes over time. The ABCD data used in this report came from https://doi.org/10.15154/z563-zd24. DOIs can be found at https://nda.nih.gov/abcd/abcd-annual-releases. The authors acknowledge the use of New Zealand eScience Infrastructure (NeSI) high-performance computing facilities and support services, funded by NeSI collaborator institutions and the Ministry of Business, Innovation & Employment. This research was supported by the New Zealand Health Research Council (Grant numbers: 21/618 and 24/838; awarded to Narun Pat), the University of Otago (awarded to Farzane Lal Khakpoor), the Neurological Foundation of New Zealand (Grant number: 2350 PRG; awarded to Jeremiah Deng and Narun Pat) and the Ministry of Business, Innovation and Employment (grants UOA2421 and RTVU2403 to Narun Pat).

## Author contributions

Khakpoor and Pat conceived and designed the study. Data acquisition, analysis and interpretation were conducted by Khakpoor, Pat, Deng and van der Vliet and Yue Wang. Khakpoor and Pat drafted the manuscript and performed statistical analyses. All authors critically revised the manuscript. Funding was acquired by Pat. Pat also supervised the project. All authors approved the final version and are accountable for the accuracy and integrity of the work. This manuscript reflects the authors’ views and does not necessarily represent those of any funding body or the ABCD Consortium.

## AI Use Disclosure Statement

During manuscript preparation, the first author used ChatGPT (GPT-5.2, OpenAI) to assist with language editing and refinement. The AI tool was used solely for editorial support. All scientific content, analyses, interpretations, and conclusions were developed by the authors.

## Availability of Data and Materials

ABCD Study data are accessible via https://nda.nih.gov/abcd upon NDA approval. All analysis code is available at https://github.com/HAM-lab-Otago-University/Ethnicity-DA

## Ethics approval statement

The ethical considerations of the Adolescent Brain Cognitive Development (ABCD) Study, including informed consent and assent procedures, participant confidentiality, and communication of assessment results, have been described in detail elsewhere (Clark et al., 2018). The Institutional Review Board (IRB) at the University of California, San Diego (UCSD; IRB #160091, approved September 13, 2016), serves as the central IRB, with reliance agreements in place across all 21 data collection sites. Prior to enrolment, written informed consent was obtained from a parent or legal guardian, and assent was obtained from all child participants.

