## Supplementaty Information for "Supervised Domain Adaptation Mitigates Cross-Ethnicity Prediction Error in Neuroimaging-Based Cognitive Prediction"

### Supplementary Information

| Single | N features | N participants | Source |
|---|---|---|---|
| Glasser Nback 2-0back | 379 | 5709 | ABCC |
| Glasser Nback 2back | 379 | 5709 | ABCC |
| Glasser Nback 0back | 379 | 5709 | ABCC |
| Glasser Nback face | 379 | 5709 | ABCC |
| Glasser Nback emotionface | 379 | 5709 | ABCC |
| Glasser Nback place | 379 | 5709 | ABCC |
| Glasser Nback face-place | 379 | 5709 | ABCC |
| Glasser Nback emotionface-neutralface | 379 | 5709 | ABCC |
| Glasser Nback NegFace-NeutFace | 379 | 5709 | ABCC |
| Glasser Nback PosFace-NeutFace | 379 | 5709 | ABCC |
| Destriuex Nback emotionface-neutralface | 167 | 7716 | ABCD |
| Destriuex Nback 2back | 167 | 7716 | ABCD |
| Destriuex Nback face-place | 167 | 7716 | ABCD |
| Destriuex Nback NegFace-NeutFace | 167 | 7716 | ABCD |
| Destriuex Nback emotionface | 167 | 7716 | ABCD |
| Destriuex Nback PosFace-NeutFace | 167 | 7716 | ABCD |
| Destriuex Nback 0back | 167 | 7716 | ABCD |
| Destriuex Nback place | 167 | 7716 | ABCD |
| Destriuex Nback 2-0back | 167 | 7716 | ABCD |
| Glasser MID Loss-Neutral | 379 | 6777 | ABCC |
| Glasser MID Reward-Neutral | 379 | 6777 | ABCC |
| Glasser MID LargeReward-Neutral | 379 | 6777 | ABCC |
| Glasser MID LargeLoss-Neutral | 379 | 6777 | ABCC |
| Glasser MID SmallLoss-Neutral | 379 | 6777 | ABCC |
| Glasser MID LossHit-LossMiss | 379 | 6777 | ABCC |
| Glasser MID SmallReward-Neutral | 379 | 6777 | ABCC |
| Glasser MID LargeReward-SmallReward | 379 | 6777 | ABCC |
| Glasser MID LargeLoss-SmallLoss | 379 | 6777 | ABCC |
| Glasser MID RewardHit-RewardMiss | 379 | 6777 | ABCC |
| Destriuex MID LargeReward-SmallReward | 167 | 9102 | ABCD |
| Destriuex MID LossHit-LossMiss | 167 | 9102 | ABCD |
| Destriuex MID Reward-Neutral | 167 | 9102 | ABCD |
| Destriuex MID SmallLoss-Neutral | 167 | 9102 | ABCD |
| Destriuex MID LargeLoss-Neutral | 167 | 9102 | ABCD |
| Destriuex MID SmallReward-Neutral | 167 | 9102 | ABCD |
| Destriuex MID LargeReward-Neutral | 167 | 9102 | ABCD |
| Destriuex MID Loss-Neutral | 167 | 9102 | ABCD |
| Destriuex MID LargeLoss-SmallLoss | 167 | 9102 | ABCD |
| Destriuex MID RewardHit-RewardMiss | 167 | 9102 | ABCD |
| Glasser SST Stop-CorrectGo | 379 | 5998 | ABCC |
| Glasser SST IncorrectStop-CorrectGo | 379 | 5998 | ABCC |
| Glasser SST IncorrectGo | 379 | 5998 | ABCC |
| Glasser SST IncorrectStop | 379 | 5998 | ABCC |
| Glasser SST IncorrectGo-CorrectGo | 379 | 5998 | ABCC |
| Glasser SST CorrectStop | 379 | 5998 | ABCC |
| Glasser SST CorrectStop-CorrectGo | 379 | 5998 | ABCC |
| Glasser SST IncorrectGo-IncorrectStop | 379 | 5998 | ABCC |
| Glasser SST CorrectGo | 379 | 5998 | ABCC |
| Glasser SST CorrectStop-IncorrectStop | 379 | 5998 | ABCC |
| Destriuex SST IncorrectGo-IncorrectStop | 167 | 8034 | ABCD |
| Destriuex SST AnyStop-CorrectStop | 167 | 8034 | ABCD |
| Destriuex SST IncorrectStop-CorrectGo | 167 | 8034 | ABCD |
| Destriuex SST IncorrectGo-CorrectGo | 167 | 8034 | ABCD |
| Destriuex SST CorrectGo-fixation | 167 | 8034 | ABCD |
| Destriuex SST CorrectStop-IncorrectStop | 167 | 8034 | ABCD |
| Destriuex SST CorrectStop-CorrectGo | 167 | 8034 | ABCD |

| | | | |
|---|---|---|---|
| Glasser Nback FC | 71631 | 4511 | ABCC |
| Glasser MID FC | 71631 | 5067 | ABCC |
| Glasser SST FC | 71631 | 4987 | ABCC |
| Glasser rest FC | 71631 | 5253 | ABCC |
| General FC | 71631 | 4273 | ABCC |
| Multitask FC | 71631 | 5027 | ABCC |
| rsfMRI subcortical-network FC | 247 | 9299 | ABCD |
| rsfMRI cortical FC | 91 | 9299 | ABCD |
| rsfMRI temporal variance | 352 | 9299 | ABCD |
| T2 gray matter avg intensity | 148 | 10471 | ABCD |
| T2 white matter avg intensity | 148 | 10471 | ABCD |
| T2 normalised intensity | 148 | 10471 | ABCD |
| T1 normalised intensity | 148 | 11177 | ABCD |
| T1 gray matter avg intensity | 148 | 11177 | ABCD |
| T1 white matter avg intensity | 148 | 11177 | ABCD |
| T2 summations | 4 | 10471 | ABCD |
| T1 summations | 9 | 11177 | ABCD |
| T1 Subcortical Volume | 19 | 11177 | ABCD |
| T2 Subcortical Volume | 19 | 10471 | ABCD |
| Total brain volume | 5 | 9141 | ABCC |
| Surface area | 148 | 9141 | ABCC |
| Sulcal depth | 148 | 11177 | ABCD |
| Cortical thickness | 148 | 9141 | ABCC |
| Cortical volume | 148 | 11177 | ABCD |
| DTI | 23 | 10194 | ABCD |

**Supplementary Figure 1** List of phenotypes.

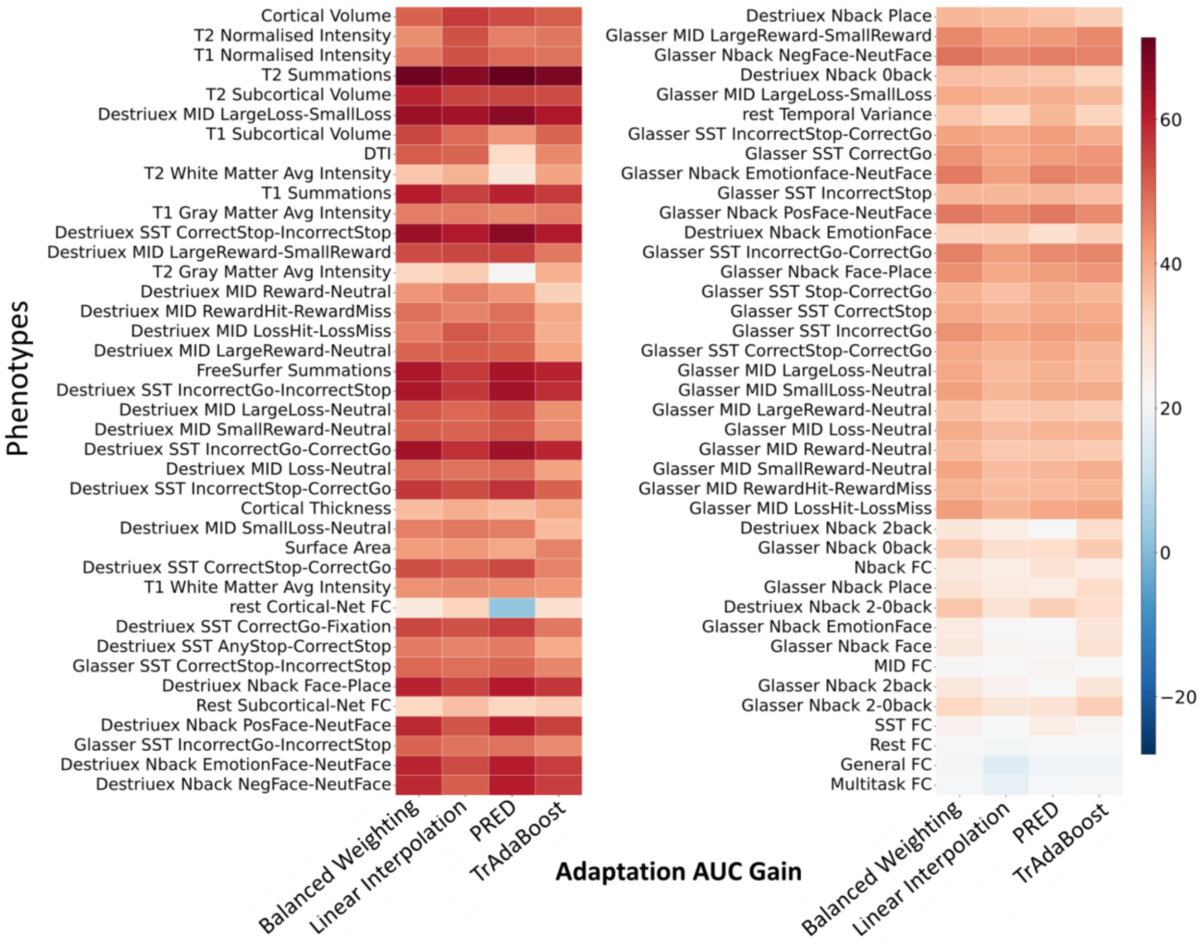


**Supplementary Figure 2** Adaptation Benefit calculated using R2 metric. The AUC gain of each adaptation method across all phenotypes sorted by their performance gap score. Starting from largest at the top left to smallest at the bottom right. A positive AUC means the adapted model produced a cumulative increase in R2 as AA labels increased.

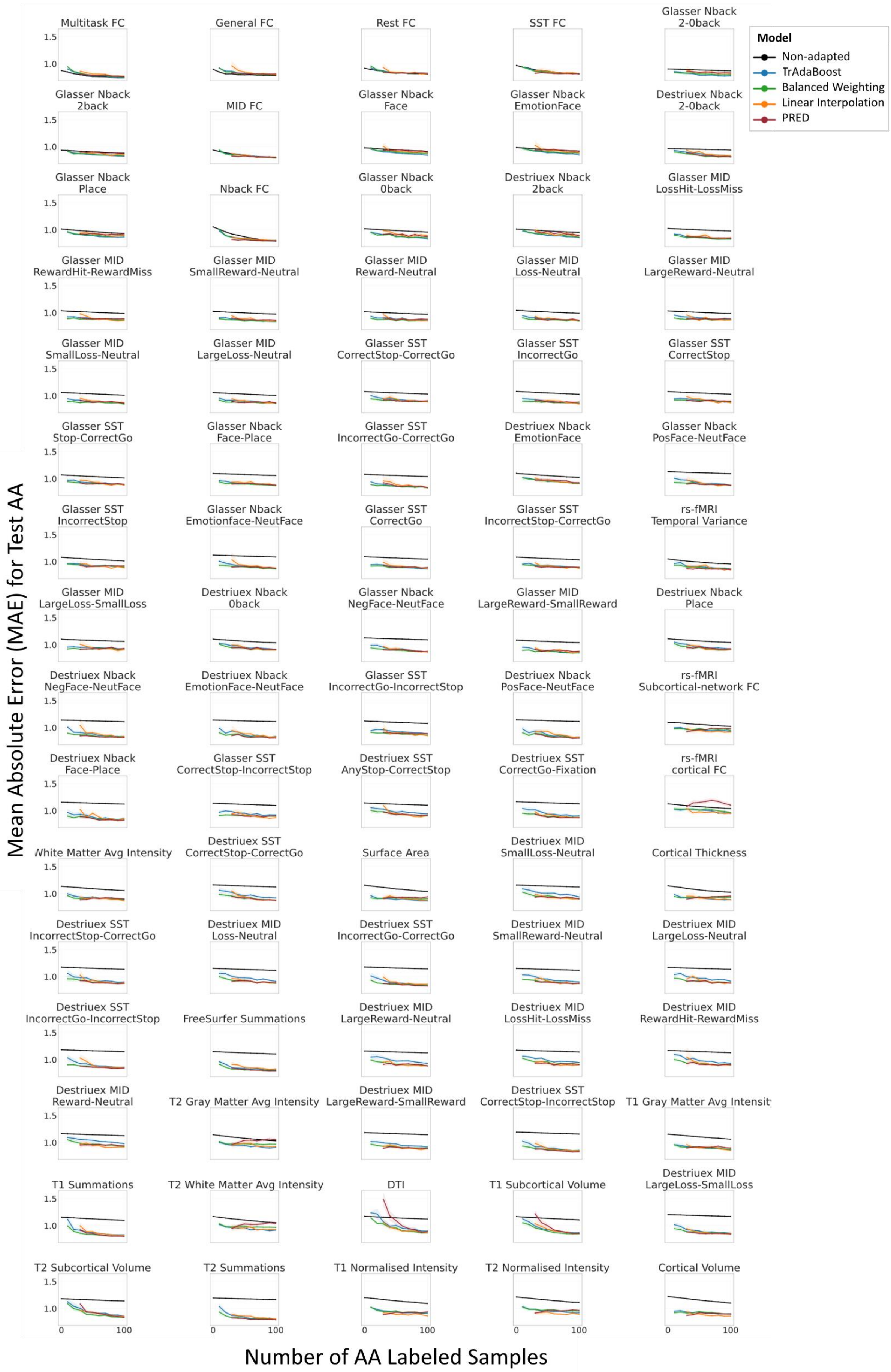


**Supplementary Figure 3** Change in AA test prediction error as increasing numbers of labelled AA participants were incorporated into model training either by direct inclusion (non-adapted) or via adaptation. Points show the mean MAE across repeated random AA subsamples at each target sample size, and shaded bands indicate approximate 95% confidence intervals across repetitions (±1.96 SD/n).
